# kmtricks: Efficient and flexible construction of Bloom filters for large sequencing data collections

**DOI:** 10.1101/2021.02.16.429304

**Authors:** Téo Lemane, Paul Medvedev, Rayan Chikhi, Pierre Peterlongo

## Abstract

When indexing large collections of short-read sequencing data, a common operation that has now been implemented in several tools (Sequence Bloom Trees and variants, BIGSI, ..) is to construct a collection of Bloom filters, one per sample. Each Bloom filter is used to represent a set of k-mers which approximates the desired set of all the non-erroneous k-mers present in the sample. However, this approximation is imperfect, especially in the case of metagenomics data. Erroneous but abundant k-mers are wrongly included, and non-erroneous but low-abundant ones are wrongly discarded. We propose `kmtricks`, a novel approach for generating Bloom filters from terabase-sized collections of sequencing data. Our main contributions are 1/ an efficient method for jointly counting k-mers across multiple samples, including a streamlined Bloom filter construction by directly counting, partitioning and sorting hashes instead of k-mers, which is approximately four times faster than state-of-the-art tools; 2/ a novel technique that takes advantage of joint counting to preserve low-abundant k-mers present in several samples, improving the recovery of non-erroneous k-mers. Our experiments highlight that this technique preserves around 8x more k-mers than the usual yet crude filtering of low-abundance k-mers in a large metagenomics dataset.

**Availability:** https://github.com/tlemane/kmtricks

**Funding:** The work was funded by IPL Inria Neuromarkers, ANR Inception (ANR-16-CONV-0005), ANR Prairie (ANR-19-P3IA-0001), ANR SeqDigger (ANR-19-CE45-0008).

## 1 Introduction

Consortia such as the 100,000 Genomes Project [1], GEUVADIS [2], MetaSub [3] and Tara Ocean [4] have generated large collections of genomic, transcriptomic, and metagenomic sequencing data, respectively. Rather than deep coverage of a single sample, such datasets contain a collection of sequencing experiments across many samples. For example, the Tara Ocean project generated metagenomic sequencing data across ecological niches all over the oceans, totalling at least 171 petabasepairs. Such valuable resources are unfortunately hard to comprehensively analyze, since their size makes bioinformatics analyses difficult.

Traditional sequence analyses such as alignment to a reference database or *de novo* assembly are both difficult and limited in the results they yield. For instance, metagenome assembly of individual samples (e.g. using MetaSPAdes [5]) is often not able to reconstruct low abundance genomes and tends to collapse variants between close strains. Co-assembly of multiple samples pools together coverage from multiple sites to alleviate this but results in further loss in strain specificity. Alternatively, aligning raw sequencing data to genome databases is hindered by the incompleteness of those databases.

A recently proposed alternative is to build an index of the raw sequencing data and then later query sequences of interest, e.g. genes or shorter sequence fragments around variants such as SNPs or indels. Traditional indexing approaches, such as those used by BLAST [6] or DIAMOND [7], do not scale to those large collections [8]. Instead, customized indexing methods have been under development. A recent review surveyed 20 tools that were all published in the last couple years, aiming to index large collections of sequencing data [8], for example `BIGSI` [9], `HowDe-SBT` [10], and `Mantis` [11]. These indexes are typically able to answer whether an arbitrary fixed-length sequence (*k*-mer) belongs to any of the samples, and, if so, which ones. Though much progress has been made, indexing a collection such as Tara Ocean has remained out of practical reach.

The vast majority of these large-scale *k*-mer indexing tools are based on common building blocks, three of them being: 1) *k*-mer counting, which summarizes each sequencing sample into a set of *k*-mers along with their abundances, 2), *k*-mer matrix construction, which aggregates lists of *k*-mer counts over a collection of samples (e.g. as in [12,13]) in the form of a *k*-mer/sample matrix with abundances as values, and 3) Bloom filters construction, where the k-mer presence/absence information for each sample is converted into a Bloom filter to save space and allow fast queries. Note that these building blocks are not specific to *k*-mer indexing tools, e.g. 1) and 3) are commonly used in short-read *de novo* assembly, and 2) also appears in transcriptomics analysis [14].

Importantly, these three steps are often categorized as “pre-processing” in *k*-mer indexing papers (e.g. [11,10]) and discounted from the running time of these indexing tools. Yet, for a dataset like Tara Oceans, these steps dwarf the running time of the subsequent index construction by up to several orders of magnitude. Although construction only needs to be done once per collection, its prohibitive running time for large collections represents an important roadblock to the usability of the tools.

In addition to the inefficiency of construction methods, sequencing errors are also dealt with sub-optimally. The work presented here is designed for indexing sequences generated by short read technologies. Even though contemporary error rates are low (0.1-0.5% as per [15]), the number of erroneous bases is large in absolute terms. Thus a vast amount of read *k*-mers contain sequencing errors and should be discarded during indexing. There are many read error-correction tools [16], that filter out *k*-mers solely by checking if their abundance is below a pre-set threshold. However, they are not a viable option for metagenomics and RNA-seq due to the presence of low-abundance genomes and the limited availability of reference genomes. Thus, current indexing methods have the unsatisfactory drawbacks of being either too conservative (discarding all low-abundant *k*-mers if the threshold is set too high), or too permissive (too many erroneous *k*-mers are kept if the threshold is set too low).

Here we propose an improved algorithm for this construction step that improves both its efficiency and the ability to discard errors. Current tools take a modular approach. They first use an off-the-shelf *k*-mer counting tool separately for each sample, and then construct a Bloom filter from the *k*-mers in that sample. We observe here that this modular approach has several drawbacks. First, it prevents fine-grain optimizations that can be obtained by tightly integrating these steps. Second, not all information is available during data structure construction, such as being able to identify all the samples to which a given *k*-mer belongs. As we will show, using such information improves the removal of erroneous *k*-mers. In summary, by limiting themselves to a modular approach, current tools leave both significant speed-up and joint filtering opportunities on the table. Given the maturity and abundance of Bloom filter-based indexing tools, as well as a plateau in performance improvement of *k*-mer counting tools [17], we believe that designing better construction algorithms through integration is an important research task.

From a collection of read samples, our method constructs Bloom filters. Based on a partitioned *k*-mer counting procedure carefully optimized for joint multi-sample counting, our novelty is the combination of three methodological ingredients, and we show that together they address the issues of long running times and sub-optimal *k*-mers filtering: 1/ We introduce a procedure for rescuing low-abundance *k*-mers at the heart of the joint multi-sample *k*-mer counting procedure. This enables more sensitive results yet discarding truly erroneous *k*-mers, saving the prohibitive indexing of all (vastly erroneous) *k*-mers. 2/ We newly apply the concept of hash counting (introduced in [18]) to the simultaneous construction of multiple Bloom filters. We partition and sort counted hashes for direct construction of Bloom filters indices without resorting to *k*-mers, saving significant time and space. In other words *k*-mers are represented by their hash value as early as possible during the entire process. 3/ We incorporate for the first time existing efficient matrix transposition techniques in a *k*-mer matrix tool, to efficiently output Bloom filter rows directly from partitioned intermediate data, saving intermediate disk space during joint multi-sample counting.

The proposed method is flexible as it offers various features depending on the user needs. Through a user-friendly pipeline, diverse results can be proposed such as count matrices instead of simple Bloom filters. Such matrices, represented in binary or plain text format, show for each row a *k*-mer and its abundance on each sample in columns. In addition to a set of linearly dependent modules along with some utilities, the tool also proposes an API for downstream developments. Rescuing low-abundance *k*-mers is also optional.

In this manuscript we restrict our attention to the Bloom filters creation pipeline using the hash counting feature and using the rescuing of low-abundance *k*-mers, both in describing the proposed method and experimenting with it. A bird-eye view of the whole pipeline showing all its potential usages is proposed in Supplementary materials, Section S4.

Using our workflow, we perform for the first time a massive-scale joint *k*-mer counting and Bloom filter construction of a 6.5 terabase metagenomics collection, in under 50 GB of memory and 38 hours, which is at least 3.8 times faster than the next best alternative.

## 2 Related works

**KMC [17] and DSK [19]** are two disk-based *k*-mer counting tools. Counting *k*-mers is an operation that identifies the set of *k*-mers present within one (or multiple) datasets and records the abundance of each *k*-mer. Intuitively, *k*-mers having a low abundance (i.e. seen few times) are more likely to be the result of one or multiple sequencing errors within reads, while *k*-mers above a certain abundance threshold are more likely to be correct. Note that neither `KMC` nor `DSK` can output Bloom filters.

In its original publication, `DSK` directly stored *k*-mers in a hash table by carefully controlling for memory and disk usage using partitioning. Recent versions of `KMC` and `DSK` are based on variants of an algorithm introduced by MSPKmerCounter [20] subsequently made popular by `KMC`. Briefly, sequencing reads are split into partitions stored on disk, then each partition is loaded in memory and *k*-mers are counted within them. Partitions are constructed in such a way that all occurrences of a *k*-mer appear within a single partition, and also, overlapping *k*-mers are attempted to be stored as longer sequences to avoid redundancy. These properties are achieved using the concepts of *minimizers* and *super-k-mers* that we will review in the Methods section, and the concept of (*k* + *x*)-mers that we will not use here.

**kmc_tools** is a binary tool included in `KMC` 3 for manipulation of its results files, which can perform e.g. set operations on list of *k*-mers, such as intersection, union, or more complex ones. However, to the best of our knowledge `kmc_tools` does not support collections of *k*-mer lists (i.e. *k*-mer matrices) and is therefore not applicable to the work presented here.

**Jellyfish [21]** is an in-memory *k*-mer counting tool that relies on an optimized hash table. One of its key advantages, besides efficiency, is that as a byproduct it constructs a *k*-mer *dictionary* : i.e. a data structure that can efficiently associate values to *k*-mers and supports efficient queries. Jellyfish was notably used to perform *k*-mer counting in `HowDe-SBT`, however its key-value store feature was not used there. Several other *k*-mer counting tools exist, and a recent benchmark [22] highlighted the peformance of `KMC3` in particular.

**Simka [23]** is a multi-sample *k*-mer counting and *k*-mer matrices construction tool that was used for metagenomics. It is based on a variation of the original `DSK` algorithm, modified to run on a distributed cluster.

**HowDe-SBT [10]** is a *k*-mer indexing method for large sequencing data collections, that extends the original concept of Sequence Bloom Trees (SBT) [24]. Briefly, `HowDe-SBT` (and in general, any SBT-based method) indexes each sample using a Bloom filter and organizes filters inside a binary tree for performing queries efficiently. `HowDe-SBT` is the most efficient variant of SBTs to date, showing fast construction and query times, yet requires an expensive pre-processing step. Precisely, the preprocessing step consists in generating from a set of samples, one Bloom filter per sample. Each Bloom filter indexes the *k*-mers considered as not erroneous contained in its corresponding set. This requires to perform a time-consuming *k*-mer counting process for each data set.

**BIGSI [9] and COBS [25]** are *k*-mer indexing methods which also use Bloom filters, however organized in a different layout than in the SBT family. These tools represent all Bloom filters of indexed samples in a flat manner, in a way that limits cache misses during query. This flat structure however does not enable to reduce redundancy between samples.

**Squeakr [18]** is a *k*-mer counting and indexing method based on Counting Quotient filters, a data structure that provides additional features compared to Bloom filters: counting, resizing, deleting and merging. Related to this work, Squeakr also counts hashes of *k*-mers.

**MetaProFi [26]** is a recent indexing scheme for *k*-mers which supports both nucleotide and amino acid sequences. `MetaProFi`, like `kmtricks`, builds a Bloom filter matrix from large sequencing data collections. However its features are different to `kmtricks` as `MetaProFi` does not apply any filtration on *k*-mers, and so, does not require to count *k*-mers.

## 3 Results

### 3.1 kmtricks A modular pipeline and library for manipulating *k*-mers from large and numerous sequencing datasets

We propose “`kmtricks`” (for “*kmer matrix tricks*”) which is a set of software components that together perform joint multi-sample *k*-mer counting, filtering, and Bloom filter construction, given as input many sequencing datasets (raw reads). Its aim is to efficiently construct Bloom filters for terabase-scale collections.

A typical execution of `kmtricks` is presented in Fig. 1. It shows the steps taken to transform reads into Bloom filters, keeping in mind that more output types (e.g. *k*-mer matrices) are supported (see Section S4). We highlight the following features which differentiates `kmtricks` from related works:

**Fig. 1.**
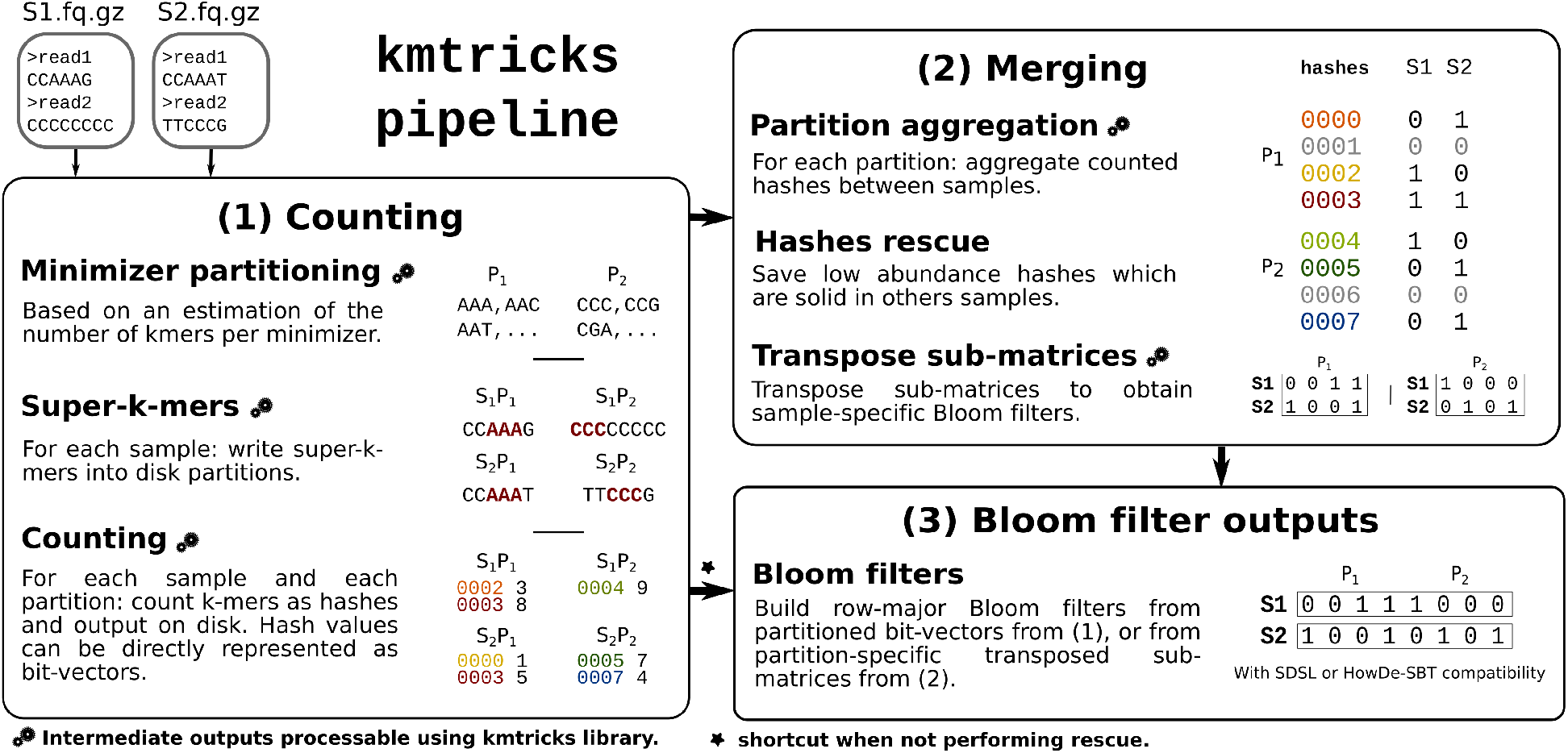
`kmtricks` pipeline overview taking as input two samples, S_1_ and S_2_. **(1) Counting:** Partitions (here, P_1_ and P_2_) over minimizers (here of length 3) are determined by sub-sampling S_1_ and S_2_ and super-*k*-mers (*k* = 5) are then written on disk according to this partitioning. Bold red sequences are minimizers (*AAA* and *CCC*). Each partition is then counted and each *k*-mer is represented by its hash value. When performing rescue, counted hashes are written on disk. Otherwise, each partition is directly represented as a bit-vector (the *⋆* symbol indicates that step (2) is skipped). **(2) Merging:** Counted hashes from equivalent partitions are aggregated and counts are binarized to produce a vector of Bloom filters (i.e. a matrix of presence/absence bit-vectors, where row indices represent hashes). This matrix is filtered using the *k*-mer rescue procedure described in section 4.3. Then, in order to build Bloom filters, i.e. having samples as matrix rows, each partition-specific sub-matrix is transposed. **(3) Bloom filter outputs:** a Bloom filter is built for each sample through concatenation of transposed submatrices (in those, each row corresponds to a sample). Bloom filters can be also obtained from first counting step if aggregation is not required. In this case, this corresponds to a concatenation of bit-vectors from (1).

— Joint *k*-mer counting allows to rescue large amounts of *k*-mers that would otherwise be discarded when processing samples independently.
— Direct counting of *k*-mer hash values instead of counting *k*-mers saves significant time for subsequent Bloom filter construction.
— `kmtricks` has been designed to be a stand-alone pipeline (Fig. 1), yet it is composed of modular tools (described in Figure S2) which are of independent interest: partitioning a set of *k*-mers (according to their minimizer), jointly count *k*-mers, construct *k*-mer matrices, transpose them, and construct Bloom filters from counted *k*-mers.
— `kmtricks` also provides a C++ library for interfacing with any stage of the pipeline, enabling for instance downstream sequence analyses based on streaming a *k*-mer matrix in row-major order.

For all presented results, the description of the data, tool versions and command lines are provided in a companion Github website (see reference [27]).

### 3.2 Scalability evaluation on RNA-seq samples

We evaluated the performance of `kmtricks` in terms of running time, peak memory usage and maximal disk space on human RNA-seq samples, while used in combination with `HowDe-SBT` for constructing indexes from Bloom filters.

The main purpose of these experiments is to show the performances of `kmtricks` in various settings (different modes, different sample sizes) and compare them to the construction of Bloom filter using a module of `HowDe-SBT` on *k*-mers counted by `Jellyfish` and `KMC3`. Results for other related tools are reported in Supplementary materials (Section S1), as they led to significantly longer run times and not used latter for large scale experiments.

These benchmarks were done on two subsets of 100 and 674 samples from a common set of 2,585 human RNA-seqs used in several *k*-mer indexing benchmarks, and first proposed in [24]. Computations were performed on the GenOuest platform on a node with 2×10-cores Xeon E5-2660 v3 2,20 GHz with 200 GB of memory. We used a SSD disk with 900 MB/s and 290 MB/s sequential read/write. All benchmarks are done using 20 cores. Results are summarized in Table 1.

**Tab. 1.**
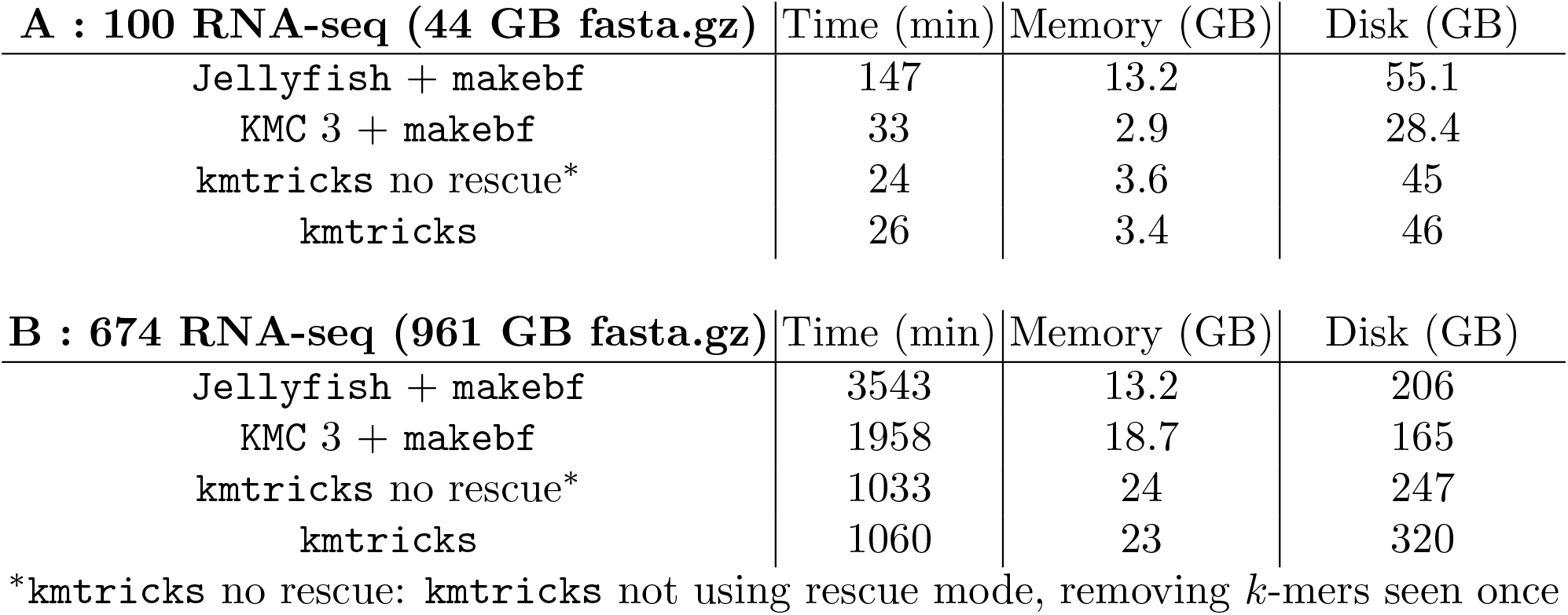
Benchmarks on two human RNA-seq datasets of respectively 100 and 674 samples. Computations were done using 20 threads with *k* = 20. Reported results correspond to Bloom filters construction (including *k*-mer counting). Memory and Disk correspond to the peak usage. `makebf` corresponds to `HowDe-SBT` sub-command which build Bloom filters from counted *k*-mers. About index constructions from Bloom filters, the performances are equivalent according to the dataset. The processes took, respectively, 21 and 120 minutes, for the 100 and 674 datasets and both used 2.6GB of memory.

| **A : 100 RNA-seq (44 GB fasta.gz)** | Time (min) | Memory (GB) | Disk (GB) |
| --- | --- | --- | --- |
| Jellyfish + makebf | 147 | 13.2 | 55.1 |
| KMC 3 + makebf | 33 | 2.9 | 28.4 |
| kmtricks no rescue* | 24 | 3.6 | 45 |
| kmtricks | 26 | 3.4 | 46 |

| **B : 674 RNA-seq (961 GB fasta.gz)** | Time (min) | Memory (GB) | Disk (GB) |
| --- | --- | --- | --- |
| Jellyfish + makebf | 3543 | 13.2 | 206 |
| KMC 3 + makebf | 1958 | 18.7 | 165 |
| kmtricks no rescue* | 1033 | 24 | 247 |
| kmtricks | 1060 | 23 | 320 |

\*kmtricks no rescue: kmtricks not using rescue mode, removing $k$ -mers seen once

On these collections, `kmtricks` is 1-1.5x faster than `KMC3` and 5-6x faster than `Jellyfish`. Thus `kmtricks` yields superior or comparable performance to other methods for indexing sequencing data, even though it performs the more complex operation of joint *k*-mer counting.

Using this dataset, we also validated that partitioning the Bloom filters does not yield uneven false positive rates that would be partition-dependent (see Supplementary Materials, Section S3).

In the next section, we show that the performance gap between other methods and `kmtricks` further widens on larger inputs, i.e. terabyte-sized collections.

### 3.3 Scaling to a large sea water metagenome collection

We performed experiments on large and complex sea water metagenomic data composed of 241 samples (distinct locations) by the Tara Ocean project [4]. This dataset is composed of approximately 6.5 thousand billion nucleotides, consisting of around 266 billion distinct *k*-mers (*k* = 20), among which 174 billions distinct *k*-mers occur twice or more (as reported by `kmtricks`).

Executions were performed on a TGCC^5^ node with 2×64-cores AMD Milan@2.45GHz (AVX2) with 512 GB of memory, on an SDD with 970 MB/s and 216 MB/s sequential read write (average on ten tests). Jobs are limited to 72h. We did not include other, non-bloom filter based, tools in this experiment because we found significantly lower performances on smaller datasets (see Supplementary Materials, Section S1).

`kmtricks` enabled to construct Bloom filters for this very large metagenomics collection in less than 24h, with similar amount of RAM and around 3.5x to 5.5x lower computation time than other methods (Table 2). Disk usage was 2x higher than other tools but 4x smaller than the compressed input data, We therefore believe that this is not a bottleneck for users. As we will see next, `kmtricks` also achieves superior results as it performs joint *k*-mer counting and is able to rescue low-abundance shared *k*-mers.

**Tab. 2.** **Comparison of construction times between kmtricks and other methods** combining *k*-mer counting with Bloom filter creation on the 6.5 terabases Tara Ocean collection using 128 threads. The `makebf` step corresponds to `howdesbt makebf` for Bloom filter creation from counted *k*-mers. The Memory and Disk columns indicate peak usage. `KMC3` and `Jellyfish` counted each sample independently and removed *k*-mers with abundance one; whereas by default, `kmtricks` performed joint *k*-mer counting and low-abundance *rescuing* (see Section 4.3) which kept some of the unit abundance *k*-mers. Results for `Jellyfish` and `KMC3` are extrapolated as our cluster jobs are limited to 72 hours. Except some rare outliers, the samples are of the same size. Computing time is therefore estimated according to the number of samples processed in 72h. For the disk usage, since `Jellyfish` and `KMC3` do single-sample counting, the peak disk usage corresponds to the Bloom filters size plus the space required to count one station. `COBS` was not executed due to its significantly longer construction times observed on smaller data and `Squeakr` is not included either because it ended in a segmentation fault on these samples.

|  | Time (min) | Memory (GB) | Disk (TB) |
| --- | --- | --- | --- |
| kmtricks | 1433 | 83.4 | 1.5 |
| Jellyfish <sup>a</sup> + makebf | $\approx 8071^b$ | $80.6^b$ | $\approx 0.8^b$ |
| KMC3 <sup>a</sup> + makebf | $\approx 5310^b$ | $100^b$ | $\approx 0.8^b$ |

<sup>a</sup>Stopped after 72h computation. <sup>b</sup>Extrapolated estimation.

As a side note, we additionally ran `HowDe-SBT` on the Bloom Filters generated by `kmtricks`, thereby creating the first complete index of all metagenomics bacterial sequences obtained in the Tara Ocean project. With Bloom filters given as input, `HowDe-SBT` ran in 1250 minutes, with a peak RAM of 165 GB. The size of the final index is 533 GB. Querying the so created index with 10,000 metagenomic reads of size 100 requires 12 minutes and 11 GB RAM.

### 3.4 Collection-aware *k*-mer filtering recovers large amounts of weak signal present in complex metagenomes

*k*-mer filtering consists in removing from a sample any *k*-mer whose number of occurrences is below a certain threshold (also called *solidity threshold*), usually set to 2 or 3. However, with data such as metagenomics or RNA-seq that have uneven coverage and include low abundance species or expressed genes, abundance does not enable to distinguish erroneous *k*-mers from real ones, as highlighted in Fig.2(a). Hence we propose to *rescue* low-abundant *k*-mers, through a *rare but shared k-mer rescue procedure*. This procedure consists in keeping any *k*-mer whose abundance is below a sample-specific threshold (called `soft-min`) whenever this *k*-mer is sufficiently abundant in one or several other samples. Section 4.3 provides a formalization and an in-depth description of the procedure.

**Fig. 2.**
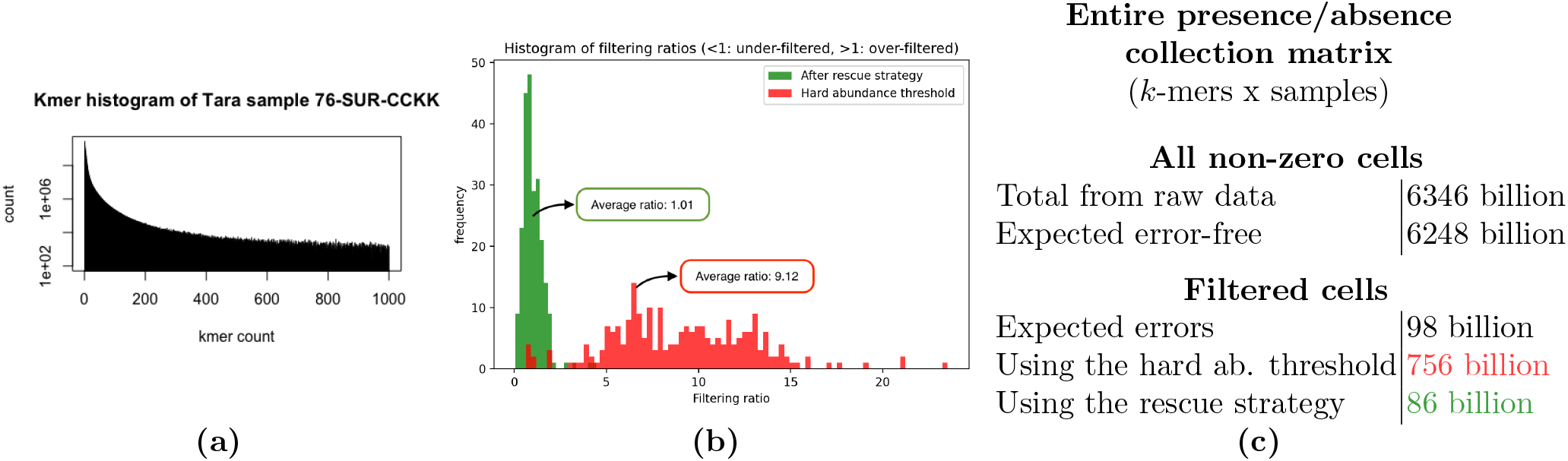
Collection-aware *k*-mer filtering recovery on Tara samples. **(a)** Kmer histogram of one of the Tara ocean samples (chosen arbitrarily), showing a flat distribution of abundances indicative of the presence of low-abundance microbes, also highlighting the lack of separation between erroneous and correct *k*-mers. **(b)** For each of the 241 Tara samples, the “Filtering ratio” reports the number of filtered *k*-mers divided by the expected number of erroneous *k*-mers (the closer to 1, the better). The green (resp. red) histogram shows the filtering ratios of samples using the `kmtricks` rescue procedure (resp. using classical removal of *k*-mers occurring only once). **(c)** Statistics of the presence/absence matrix (*k*-mer x sample) for the entire Tara collection. The “All non-zero cells” table concerns the total number of cells in the matrix where a *k*-mer is present in a dataset (non-zero abundance), and the “Filtered cells” table reports the total estimated number of erroneous *k*-mers inside each dataset (“Expected errors”), and the number of cells filtered out by the absolute threshold strategy and the rescue strategy.

We compared an usual filtering method, i.e. discarding the *k*-mers seen once (`hard-min` = 2), with our rescue strategy applied on the whole *k*-mer spectrum (`hard-min` = 1). We validated this strategy as follows. For each sample we computed

1. *err*_*th*_: the theoretical expected number of erroneous *k*-mers,
2. *err*_*one*_: the number of *k*-mers occurring only once, and
3. *err*_*unrescued*_: the number of *k*-mers that are still considered as erroneous after our rescue procedure.

We then look at the ratio *err*_*unrescued*_*/err*_*th*_ and compare it to the ratio *err*_*one*_*/err*_*th*_ that would be obtained by a classical filtering of *k*-mers occurring once. The closer a ratio is to one, the better.

The Tara Oceans dataset was mainly generated by HiSeq 2000 technology (222 samples out of 241), 8 samples were generated by HiSeq2500, 4 samples by GAIIx. For each of these technologies, we computed the theoretical error rate *err*_*th*_. Given raw sequencing data from *Acinetobacter baylyi* generated by these three sequencing technologies ^6^ we counted the number of erroneous *k*-mers (*k* = 20), i.e. those absent from the reference genome. Our estimated *k*-mer error rates (*e*.*g*. ratio of erroneous *k*-mers) are respectively 1.27%, 9.27%, and 3.38% for HiSeq2000, HiSeq2500 and GAIIx. As a side note, from those ratio, one can estimate respective base error error rates being respectively 0.06%, 0.48%, and 0.171%.

Results shown in Fig.2(b) and (c) highlight the importance and efficacy of our *k*-mer rescue procedure. Indeed, the quantity of *k*-mers filtered out is close to the theoretical expected value when using the rescue procedure (average ratio of 1.01, 86 billions *k*-mers filtered compared to 98 billions expected to be filtered). Around 8-9x too many *k*-mers appear to be wrongly filtered out when removing from each sample *k*-mers occurring only once (756 billions *k*-mers filtered compared to 98 billions expected to be filtered, and average filtering ratio of 9.12).

Thus, when indexing datasets containing low-abundant sequences as in the Tara ocean metagenomes, *k*-mer rescuing appears to be essential as it: **1**. side-steps the issue of removing low-abundant *k*-mers which ends up discarding an order of magnitude too many *k*-mers; **2**. recovers a number of *k*-mers close to the expected one using co-occurrence across samples.

## 4 Methods

### 4.1 Definitions

A *minimizer* of length *m* within a sequence *s* is the smallest *m*-mer within *s*, where typically “smallest” is understood in the lexicographical sense. A *super-k-mer* is a sequence in which all constituent *k*-mers have the same minimizer.

A *Bloom filter* [28] is an approximate membership query (AMQ) data structure that allows two operations: inserting and querying elements *u* ∈ 𝒰. It is a bit array *B*[0..*n*] with *l* hash functions *h*_*i*_ : 𝒰 → {0, …, *n*} ∀*i* ∈ [1..*l*]. Insertion can be defined as follows *B*[*h*_*i*_(*x*)] ← 1, ∀*i* ∈ [1..*l*] and lookup as 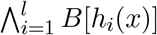. Lookups can return false positives but no false negatives. In the following, we will consider Bloom filters with a single hash function (*l* = 1).

We use the terms *color-aggregative* and *k-mer-aggregative* as defined in [8]. A color-aggregative data structure represents within a single index all *k*-mers of the collection, and each *k*-mer is associated to its pattern of presence/absence across the whole collection. Conversely, a *k*-mer-aggregative data structure constructs separate *k*-mer indices, one per sample.

The strand of each sequenced read being unknown, in `kmtricks`, as in all *k*-mer counting and indexing tools, each *k*-mer is represented by its canonical representation: the smallest string (e.g. in the lexicographic order) between itself and its reverse complement.

We refer to *hash counting* as the process of counting hash values of a set of elements instead of counting the elements (*k*-mers) themselves. This is the counterpart of *k*-mer counting except that here *k*-mers are converted into (non-invertible) hash values, and several *k*-mers may collide to the same hash value. Classical *k*-mer counters (e.g. `Jellyfish`, `KMC`) also use hash values inside their algorithms, but always return exact *k*-mers in the final results. Here ‘hash counting’ goes one step further and discards the original *k*-mer sequence, as it is unnecessary for constructing Bloom filters. The key difference between performing hash counting and directly constructing a Bloom filter is that we record the abundances of *k*-mers, which is essential for further filtering. This is akin to Count-Min sketches [29] however we focus on the efficient construction of an external memory representation which is not meant to support random accesses.

Previously, `Squeakr` performed hash counting within Counting Quotient filters. This method however cannot directly be applied to efficiently construct Bloom filters, as we do in this work by counting, partitioning and sorting hashes prior to data structure construction. In [18], hash counting is applied to construct each CQF at a time, later merged using Mantis [11]. In `kmtricks`, we directly construct multiple Bloom filters by partitioning counted hashes, saving time.

### 4.2 A modular pipeline for large-scale Bloom Filters construction: kmtricks

`kmtricks` supports the construction of either a *k*-mer matrix or Bloom filters. In both cases, the input is a collection of sequencing data files in FASTA/FASTQ format. The output is either a matrix having *k*-mers or hashes as rows, samples as columns, and *k*-mer counts as values, or a collection of Bloom filters, one per sample. In the following, for simplifying the reading of this manuscript, we focus on the process of constructing Bloom filters, counting hash values instead of *k*-mers. A general presentation of the whole pipeline with all its features is proposed in Supplementary materials, Section S4.

In other tools, the construction process of Bloom filters can typically be broken into two steps: 1) efficiently counting *k*-mers then 2) inserting distinct *k*-mers into filters, on a per-sample basis. `kmtricks` streamlines this process by realizing that in the case of Bloom filters, only the hashes need to be counted, not *k*-mers. Furthermore, in order to cope with terabytes of input data and still be able to efficiently count hashes, a careful partition-aware hashing scheme is designed.

#### 4.2.1 Partitioning

`kmtricks` performs parallel *k*-mer counting following the classical paradigm of partitioning *k*-mers based on their minimizers and then constructing super-*k*-mers, as introduced by `KMC2` [17]. However, the process is newly modularized so that intermediate tasks correspond to separate programs. Conceptually, the set of all possible minimizers is first partitioned in the following balanced way: all partitions should contain a roughly equal total number of *k*-mers. This is performed by the GATB library [30] that implements the same algorithm as in `DSK` [19] (and `KMC` 2 & 3 [31]).

In order to take advantage of the partitioning scheme, we use *partitioned Bloom filters* (pBFs). These are Bloom filters that are partitioned into *P* sub-filters with exclusive (and consecutive) hash spaces *h*_*p*_ : 𝒰_*p*_ → {*p* × *s*, …, *p* × *s* + *s* − 1} with *p* ∈ [0..*P* − 1] and 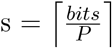 (rounded up to a multiple of 8) with “*bits*” corresponding to the user-requested Bloom filter size. pBFs allow to populate only a small part of a Bloom filter when processing a *k*-mer partition, which saves memory and enables coarse-grained parallelization at both the construction and query stages. A classical Bloom filter can be obtained by a simple concatenation of the pBFs thanks to the consecutive hash spaces. This entails to adapt the query operations, see Section 4.2.5.

#### 4.2.2 Counting

This step consists in computing for each sample its super-*k*-mers and writing them in their corresponding partitions on disk (Fig 3.a). From those super-*k*-mers, *k*-mer hash values are de-duplicated and the abundance of each distinct hash value is determined within each partition.

**Fig. 3.**
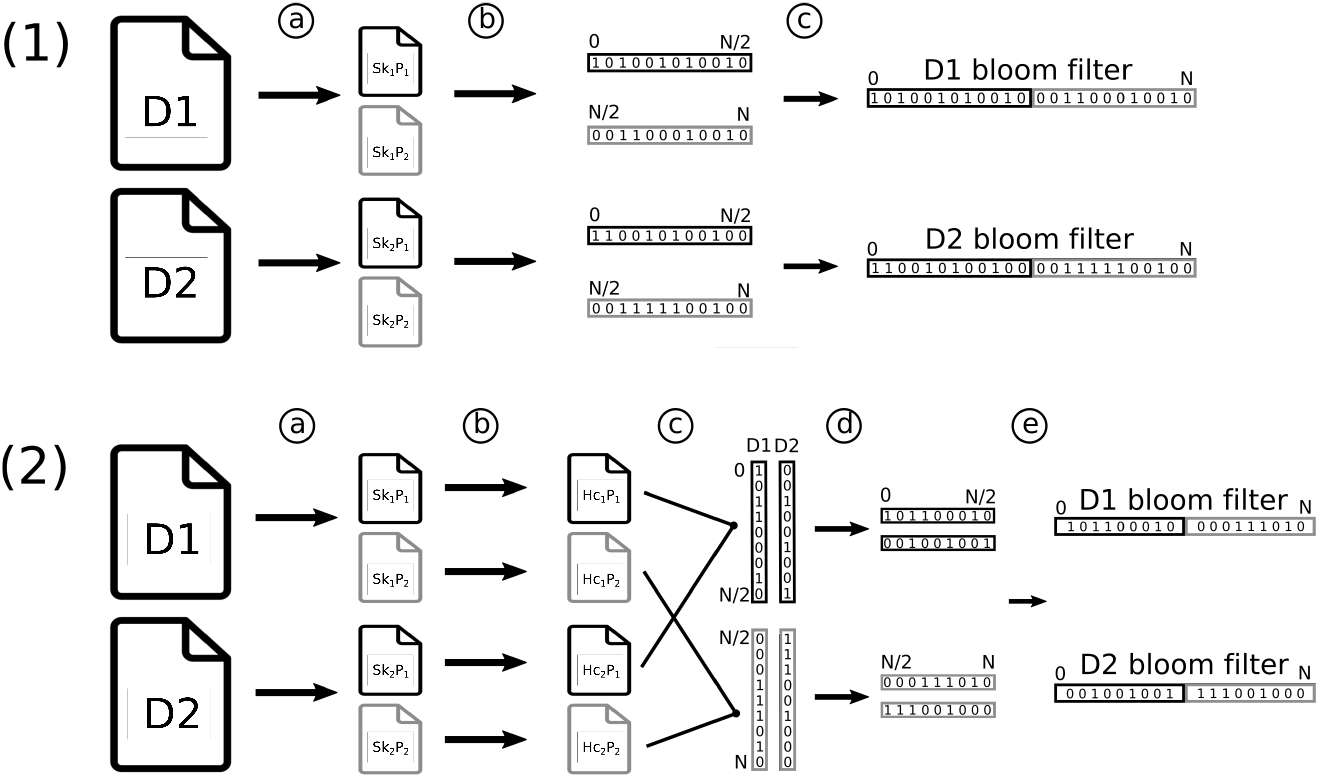
Bloom filters construction pipeline with two samples D1 and D2 using two partitions: black (1) and gray (2). **Sk** and **Hc** denote respectively super-*k*-mers and hash counted **(1) Bloom filters pipeline without** *k***-mer rescue**: 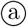 Divide sample into partitioned super-*k*-mers. 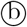 Split super-*k*-mers into *k*-mers before hashing them and counting hashes in partitions. For each partition, output presence/absence bit-vectors, i.e. partitioned Bloom filters. 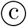 Concatenate equivalent partitions between samples to obtain one Bloom filter per sample. **(2) Bloom filters pipeline with rare** *k***-mer rescue**: 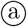 same as (1). 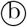 same as (1), but output hashes and their counts. 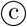 Merge and binarize (according to the rescue procedure, see 4.3) equivalent partitions to build one sub-matrix per partition with pBFs in columns. 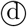 Transpose sub-matrices to obtain pBFs in rows. 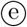 same as (1)-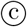.

`kmtricks` optimizes the output in the following way. If *k*-mer rescue (see 3.4) is not performed, bit-vectors are output immediately instead of a list of counted hashes (Fig 3.1.b). One bit vector is output per partition and per sample. In other words, these bit-vectors correspond to pBFs built from the hashes.

Otherwise if *k*-mer rescue is performed, bit-vectors cannot be immediately output as counts of the same hash value must be examined over all samples. In this case, hashes and their counts are dumped to disk for each partition from each sample (Fig. 3.2.b).

#### 4.2.3 Merging in *k*-mer rescue mode

If *k*-mer rescue is performed, partitions of hash values need to undergo a merging step in order to obtain a collection of Bloom filters. Aggregation of hashes over multiple samples is achieved using the classical *n*-way merge algorithm on equivalent partitions across samples. This algorithm assumes sorted inputs, yet the counting step already provides a sorted output.

The count vector of each row (corresponding to a single hash value) is processed according to the *k*-mer rescue procedure, details are given in Section 4.3. Row count vectors are transformed into a binary representation during the merge step. In a partition, all possible hashes are considered. This means that for each missing hash value (corresponding to *k*-mer(s) not seen in the partition), an empty bit-vector is appended to the matrix. Hashes are not stored (only the bit-vectors are), as they implicitly correspond to row indices. At the end of this step, we have *P* sub-matrices of 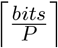 presence/absence bit-vectors each.

At this point, the resulting matrices are color-aggregative, i.e. each row represents the presence or the absence of the corresponding hash value across samples. One seeks to convert the data into a *k*-mer-aggregative representation, i.e. where each filter represents hashes for a single collection. Switching from a color-aggregative representation to a *k*-mer-aggregative representation can be achieved through a bit-matrix transposition. The matrices are additionally sorted so that each bit-vector row corresponds to consecutive hashes in {*p* × *s*, …, *p* × *s* + *s* − 1}.

When performing a transposition, we transform a matrix with hashes in rows associated with presence/absence bit-vectors into a matrix with samples in rows associated with a one-hash pBF. Due to Bloom filter partitioning, *P* transposed matrices are in fact obtained, each with a number of rows corresponding to the number of samples. The horizontal concatenation of each corresponding row from these matrices allows one to build one Bloom filter per sample.

#### 4.2.4 Outputs

`kmtricks` can produce different outputs as described in Figure S2. In the context of Bloom filter construction, it produces hashes presence/absence vectors (kmer-aggregative) or pBFs vectors (color-aggregative) both seen as bit matrices. Subsequently, pBFs vectors can be converted into sample-specific Bloom filters that are compatible with the SDSL library [32] and `HowDe-SBT`.

#### 4.2.5 Query

Bloom filters can be queried in the usual way, apart from a small technicality: `kmtricks` creates Bloom filters that are partitioned according to minimizers. The minimizer of each *k*-mer from a query sequence must be computed in order select the correct hash function. To facilitate the queries, we propose an `HowDe-SBT` wrapper compatible with our hashing scheme, through the `kmtricks query` command. The `kmtricks` C++ library can additionally be used for directly querying the Bloom filters in the correct manner.

### 4.3 A novel technique to rescue rare *k*-mers

To rescue low-abundant but likely correct *k*-mers, as performed in Section 3.4, we design a rather simple technique based on examining the abundance of each *k*-mer across sequencing samples. This technique is only practically applicable in conjunction with joint *k*-mer counting. It cannot be directly implemented in a one-sample-at-a-time construction procedure, unless such procedure discards no *k*-mer which would result in prohibitively large intermediate storage. When a low-abundant *k*-mer is observed in a sample (with abundance lower than a user-defined threshold), its abundance in other samples is used to decide whether to keep its abundance for that sample or not.

In each sample *i*, two thresholds are used: *k*-mers whose abundance is below a first threshold called `hard-min`_*i*_ are simply discarded from sample *i* during the counting step. Such *k*-mers cannot be rescued latter. Among remaining *k*-mers, those whose abundance is higher than or equal to a second threshold called `soft-min`_*i*_ (with `soft-min`_*i*_ ≥ `hard-min`_*i*_) are said to be “solid” and are conserved during the merge step. Finally a *k*-mer whose abundance in sample *i* is in [`hard-min`_*i*_, `soft-min`_*i*_[is conserved only if there exists at least `share-min` other samples in which this *k*-mer is solid. The parameter `share-min` is user-defined and is independent from the considered sample.

The thresholds `hard-min`_*i*_ and `soft-min`_*i*_ are user-defined and can be set independently for each sample *i*, or a unique value can be set to all samples. To facilitate usage, `soft-min`_*i*_ can be automatically determined with respect to the total number of *k*-mers in set *i*. In this case `soft-min`_*i*_ is the smallest value ≥ 1 such that the number of *k*-mers occurring `soft-min`_*i*_ is smaller than a user-defined percentage of the total number of *k*-mers.

Figure 4 presents several examples showing the application of those thresholds.

**Fig. 4.**
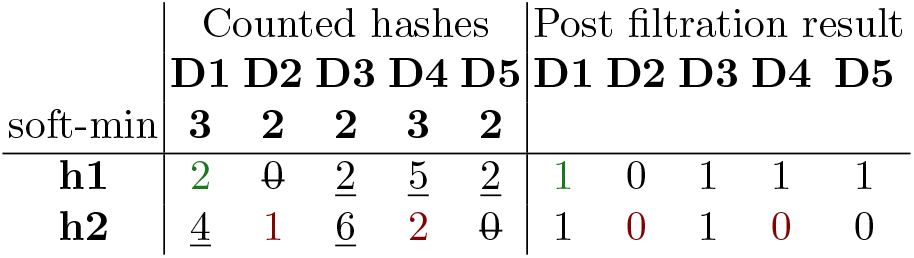
Rescue procedure example for two hash values (h1 and h2) and five samples (D1 to D5 using sample-specific `soft-min` and the following set of parameters: `hard-min=1` (for all samples), and `share-min=3`. The strikeout values are lower than `hard-min`, underlined values are higher or equal to `soft-min` in their sample (solid *k*-mer). Other values can be rescued. Among them the value shown in green is rescued, as **h1** has a abundance lower than 3 in D1 but it is solid in at least `share-min` other samples (D3, D4, D5). Values shown in red are not rescued as **h2** is solid only in 2 samples.

## 5 Discussion

We propose `kmtricks`, a novel method for efficiently counting *k*-mers across multiple samples and for generating Bloom filters. In addition to being the fastest method for generating Bloom filters over terabyte-scale collections, our approach `kmtricks` proposes a novel mechanism to filter erroneous *k*-mers using their co-occurrence across samples, i.e. going beyond filtering on a per-sample basis. This approach leads to significantly improved recovery of *k*-mers in metagenomes. `kmtricks` is flexible as, in addition to this adjustable possibility for subtle *k*-mer filtration, it offers diverse usage scenario: Bloom filters creation, *k*-mer matrices and various operations on each module output. The scope of `kmtricks` is therefore Bloom filter construction, k-mer matrix construction, but it is not a drop-in replacement for k-mer counting tools.

In our tests on relatively small collections (100-674 RNA-seq datasets, with on average hundreds of millions of distinct *k*-mers per sample) the performance of `kmtricks` is roughly equivalent to the state of the art `KMC` 3 *k*-mer counter combined with the Bloom filters construction procedure of `HowDe-SBT`. However `kmtricks` stands out on larger collections having higher number of *k*-mers per sample, such as Tara Ocean (*>* 6 TB of sequences, with several billions of distinct *k*-mers per sample). For those, Bloom filter construction becomes a bottleneck and highlights the superior efficiency of the streamlined `kmtricks` pipeline.

At a high level, `kmtricks` is able to output matrices either in column-major order or in row-major order, where rows can either be *k*-mers or hash values. This flexibility allows `kmtricks` to provide inputs for both families of indexing data structures (*k*-mer-aggregative and color-aggregative, as defined in [8] and recalled in the Methods section). Row-major order makes the presence/absence of a *k*-mer directly accessible across all samples, and is contiguous in memory. In column-major order, each Bloom filter is independent and provides information about the existence of all *k*-mers in one sample.

`kmtricks` uses a particular hash function on *k*-mers that partitions the hash space by minimizers. We experimentally validated that this scheme does not introduce more false positives than a classical Bloom filter.

Contemporary of `kmtricks`, MetaGraph [33] has been very recently proposed as an exact *k*-mer membership indexing scheme (i.e. not using Bloom filters). MetaGraph has been applied to collections of hundreds of terabases using cloud resources. Its preprocessing step is `KMC` 3, and it does not perform *k*-mer rescue. Therefore its features are different to `kmtricks`, yet we added a performance comparison in Table 1. Ideally, `kmtricks` could be integrated within a MetaGraph-like approach to combine single-server efficiency and *k*-mer rescue within a cloud architecture.

## Acknowledgements

This work used HPC resources from the TGCC of CEA and the GenOuest bioinformatics core facility (https://www.genouest.org). The authors are grateful to Bob Harris for discussion on `HowDe-SBT`, and Eric Pelletier & Jean-Marc Aury who provided links to Tara and *Acinetobacter* datasets, and precious information about these data.

## Supplementary

### S1 Human RNA-seq benchmarks

This section present an extensive version of the Table 1 (Section 3.2) with additional comparisons against non-Bloom filter based tools. Tool versions are shown in Table S4. Details about data and scripts are available from the `kmtricks` github companion website (see reference [27]). A Conda environment is also provided to reproduce these benchmarks.

**Tab. S1.** Benchmarks on two human RNA-seq datasets of respectively 100 and 674 samples. Computations were done using 20 threads with *k* = 20. However as `COBS` supports only `McCortex`-file for *k* = 31, we also propose results for `kmtricks` + `HowDe-SBT` using *k* = 31. For Time and Memory, when two values are provided in a cell, the first corresponds to the pre-processing time (*k*-mer counting and possibly Bloom filter creation) and the second to the index construction. Memory and Disk correspond to the peak usage. Disk usage corresponds to the total required space to build the index, including temporary files, Bloom filters and the final index. For `McCortex`-`COBS`, the disk usage corresponds mainly to the ctx files from `McCortex`.

| kmer counter (& bf creation) | Index | Time (min) | Memory (GB) | Disk (GB) |
| --- | --- | --- | --- | --- |
| **A : 100 RNA-seq (44 GB fasta.gz)** |  |  |  |  |
| Jellyfish (& makebf) | HowDe-SBT | 147 + 21 | 13.2 — 2.6 | 55.1 |
| KMC 3 (& makebf) | HowDe-SBT | 33 + 21 | 2.9 — 2.6 | 28.4 |
| McCortex $k = 31$ | COBS | 256 + 67 | 27 — 1.5 | 327 |
| Squeakr | Mantis | 64 + 24 | 3.6 — 27.8 | 25.8 |
| kmtricks | HowDe-SBT | 24 + 21 | 3.6 — 2.6 | 45 |
| kmtricks<sup>R</sup> | HowDe-SBT | 26 + 21 | 3.4 — 2.6 | 46 |
| kmtricks<sup>R</sup> $k = 31$ | HowDe-SBT | 20 + 21 | 3.6 — 2.6 | 50 |
| **B : 674 RNA-seq (961 GB fasta.gz)** |  |  |  |  |
| Jellyfish (& makebf) | ∅ | 3543 | 13.2 | 206 |
| KMC 3 (& makebf) | ∅ | 1958 | 18.7 | 165 |
| KMC 3 | Metagraph | 561 + 973 | 18.7 — 21.9 | 96.3 |
| kmtricks | ∅ | 1033 | 24 | 247 |
| kmtricks<sup>R</sup> | HowDe-SBT | 1060 + 120 | 23 — 2.4 | 320 |

kmtricks<sup>R</sup>: kmtricks using rescue mode

#### Results not including the index creation

As shown Table ±S1, on the smaller dataset (100 RNA-seq, 44 GB fasta.gz), `kmtricks` outperformed `Jellyfish` used in combination with `makebf`, `McCortex` and `Squeaker` in term of computing time (by 2.6-10x) and memory usage (by 1-3.9x) and use comparable disk space. We also substituted `Jellyfish` with `KMC3` in `HowDe-SBT`, yielding comparable time/memory performance to `kmtricks` on this collection. In terms of *k*-mer counting alone, `KMC3` is 1.8x faster with similar memory usage, however `KMC3` does not create Bloom filters from counted *k*-mers, and does not support joint *k*-mer counting and so can not provide a similar *k*-mer rescue procedure. Its integration in a Bloom filter construction pipeline turns out to be significantly less scalable than `kmtricks` as shown Section 3.3, dealing with larger and more complex data.

One the larger dataset (674 RNA-seq 961 GB fasta.gz), similar conclusions hold, `kmtricks` remaining the fastest tool to provide Bloom filters from raw read files (1.8-3.3x faster).

#### Results including the index creation

We used `HowDe-SBT` from Bloom filters and `COBS` and `Mantis` from counted *k*-mers for constructing final indexes. Except for `COBS` which is significantly longer than other tools (3.2 times longer than `HowDe-SBT`) performances are equivalent.

Even if it is currently not published, we also tested `Metagraph` (https://github.com/ratschlab/metagraph) on the largest dataset and using `KMC3` as prepossessing step. Compared to `HowDe-SBT` using `kmtricks` as a prepossessing step, and including the rescue mode, `KMC3` +`Metragraph` uses 3.3 times less disk, and is 1.3 times slower, while using slightly less RAM (21.9 GB versus 23 GB).

#### k-mer matrix construction

In this manuscript, we focused on the Boom filters construction. However, kmtricks is able to build different type of matrices like abundance or presence/absence matrices. In the table S2, we present a quick comparison between Bloom and abundance matrix construction. Since these two modes share a part of their algorithms, their performances are often close in terms of computing cost. This can of course differ depending on the parameters such as very large Bloom filter size for instance. Moreover, in Bloom mode, the fixed size of hashes allows us to use a more efficient compression algorithm, in both time and space, resulting in a less-intensive IO usage.

**Tab. S2.** Comparison of *k*-mer matrix and Bloom filter matrix construction on 100 RNA-seq samples. Computations were done using 20 threads and *k* = 20.

| 100 RNA-seq (44 GB fasta.gz) |  |  |  |
| --- | --- | --- | --- |
|  | Time (min) | Memory (GB) | Disk (GB) |
| **kmtricks** Bloom | 26 | 3.4 | 46 |
| **kmtricks** *k*-mer | 29 | 2.8 | 54 |

#### k-mer counting

Although `kmtricks` is not a drop-in replacement for *k*-mer counters, we compared it with `Jellyfish` and `KMC3` on 100 RNA-seq samples. The results are presented in the table S3. As shown in the table, `kmtricks` is faster than `Jellyfish` but it should be noted that the outputs are different since `Jellyfish` produces a hash table. For the comparison with `KMC3`, the performances are close because `kmtricks` is adapted to multi-sample counting. For single-sample counting, a *k*-mer counter like `KMC3` is probably more adapted and efficient.

**Tab. S3.** Comparison of *k*-mer counting on 1 and 100 RNA-seq samples. Computations were done using 20 threads and *k* = 20.

| 100 RNA-seq (44 GB fasta.gz) |  |  |  |
| --- | --- | --- | --- |
|  | Time (min) | Memory (GB) | Disk (GB) |
| **kmtricks** | 26 | 2.2 | 48 |
| **Jellyfish** | 51 | 6.2 | 49 |
| **KMC3** | 24 | 2.9 | 29.8 |

### S2 Tool versions

**Tab. S4.** Tool versions.

| Tool | Version or git sha1 |
| --- | --- |
| **HowDe-SBT** | 2.00.02 |
| **Jellyfish** | 2.3.0 |
| **KMC** | 3.1.1 |
| **McCortex** | 1.0.1 |
| **COBS** | 1915fc0 |
| **Squeakr** | 0.6 |
| **Mantis** | 0.2.0 |
| **Metagraph** | 0.1.0 |
| **kmtricks** | 1.1.1 |

### S3 Empirical analysis of pBFs false positive rate

Since `kmtricks` Bloom filters are partitioned (pBFs), a potential drawback is that the partition repartition is uneven and that false positive rate is partition-dependant. We checked the false positive rate of each partition and performed the following experiment: given a pBF of total size *s*, we compared for each of its partitions the actual false positive rate versus the false positive rate that would be obtained by a non-partitioned Bloom filter of size *s* (called the theoretical false positive rate). We computed the partition-dependent false positive rate (using 300 partitions) for a dataset with 100 human RNA-seq samples. Results shown in Fig. S1 give the false positive rate dispersion across partitions for 15 samples compared to the theoretical false positive rate of these 15 samples. Results on the remaining 85 samples are similar. Command lines and full results are available at github.com/pierrepeterlongo/kmtricks_benchmarks. Despite some outliers, partition-dependent false positive rates remain close to the theoretical values.

As the partitioning scheme is the same for all samples of a dataset, it is theoretically possible for some experiments (very heterogeneous for instance) and some samples that the false positive rate variation across the partitions is more important than what we observe here. For allowing query-time correction of this effect, `kmtricks` provides as an output the false positive rate of each partition for each sample.

**Fig. S1.**
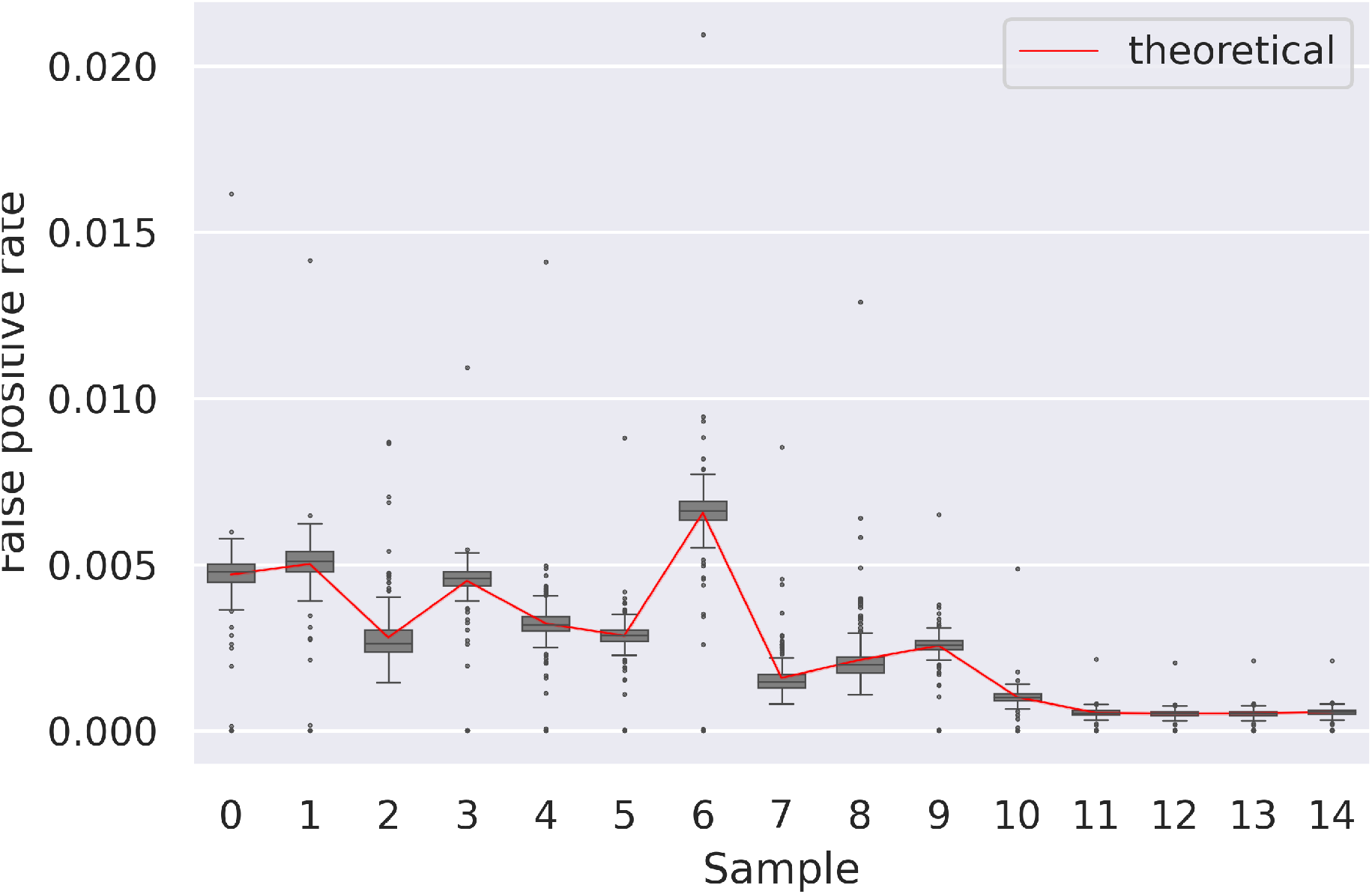
Partition-dependant pBF false positive rate. Given 15 human RNA-seq samples, the distribution of false positive rates across partitions is shown as well as the theoretical false positive rate, obtained with no partitioning.

### S4 kmtricks modules

**Fig. S2.**
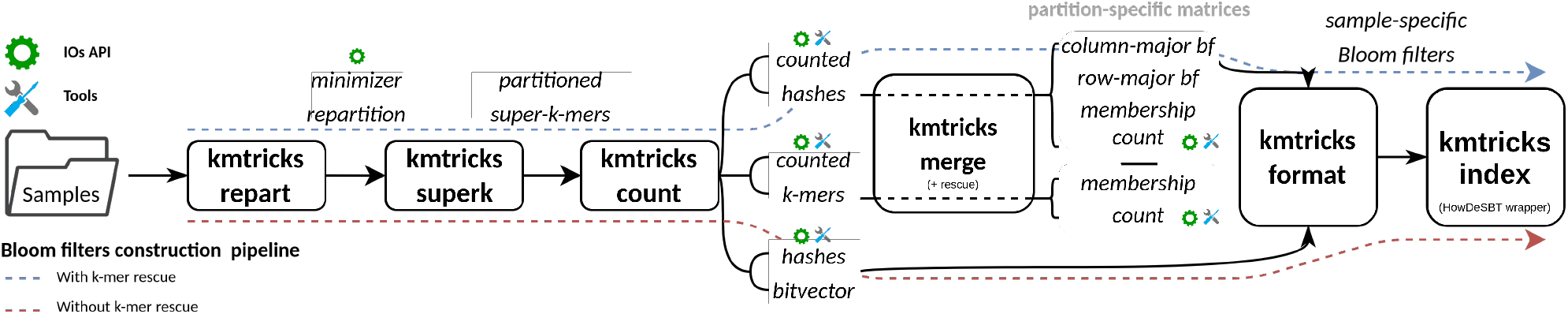
`kmtricks` modules overview. The different possible paths in `kmtricks`’s pipeline are represented by a diagram of modules (boxes) annotated with their intermediate outputs (italics). Many of the intermediate outputs are readable by the `kmtricks` API, and tools are also available for basic operations such as `dump` or `aggregate`. The two dotted lines show the pipeline described in the paper, i.e. Bloom filters construction with (blue) and without (red) *k*-mer rescue.

`kmtricks` tool suite is composed of a set of linearly dependent modules along with some utilities and API allowing *k*-mer/hash/bf matrices construction. As described in Figure S2, each module corresponds to one step of the `kmtricks pipeline` but some can have different inputs/outputs depending on the chosen output mode (*k*-mer or hash counting, with or without *k*-mer rescue, etc…).

Additional modules are provided to exploit `kmtricks`’s files: 1) `kmtricks dump`, allowing to convert individual files in human readable format. 2) `kmtricks aggregate`, allowing to aggregated consistent files, e.g. all count sub-matrices or all counted partitions of one sample. In the same spirit, an API provides sequential reading of `kmtricks`’s files allowing for instance parallel streaming of *k*-mer matrices from counted *k*-mer partitions.

## Footnotes

5. Trés Grand Centre de Calcul of CEA (http://www-hpc.cea.fr/fr/complexe/tgcc.htm)

6. Jean-Marc Aury, Genoscope, personal communication.

